# An Investigation of mini-GRID Radiation therapy with Immune Checkpoint Blockade in a Murine Tumor Model

**DOI:** 10.1101/2025.09.14.676104

**Authors:** Patrick Sansone, Ashlyn G Rickard, Nerissa T Williams, Rico J Castillo, Simon Brundage, Yvonne M Mowery, Mark Oldham

## Abstract

Spatially fractionated radiotherapy has shown potential to improve therapeutic outcomes possibly with an immunogenic mechanistic component. Here we report on *in vivo* mouse studies investigating mini-GRID pencil-beam radiotherapy combined with anti-PD-1 immune checkpoint blockade. Methods: GRID therapy was delivered at 225kV using the XStrahl Small Animal Radiation Research Platform with two custom lead mini-GRIDs, each consisting of an array of equally spaced holes: 1 mm diameter with 1mm spacing and 254 µm diameter with 508 µm spacing. GRID dosimetry was characterized using EBT3 film to determine peak-to-valley dose ratios and output. Two studies were performed with C57BL/6J mice bearing subcutaneous LLC1 flank tumors. In the first, mice (n=5/group) were treated in 3 groups with a single fraction: 15 Gy open field, 15 Gy 1 mm GRID, or 24 Gy 1 mm GRID. In the second, mice (n=6-7/group) were treated with fractionated GRID radiotherapy in 5 groups: 15 Gy open field x 3 fractions, 15 Gy hemi-irradiation x 3 fractions, (15 Gy 1 mm GRID x 3 fractions, or 15 Gy 254 µm GRID x 3 fractions. All mice were treated with 200 μg anti-PD-1 antibody on days 0, 3, and 6, then weekly until humane endpoint (tumor >15 mm in any dimension or ulceration). Results: Peak to valley ratios were 24.5 ± 0.6 and 19.8 ± 0.7 for the 1 mm and 254 µm GRIDs, respectively. Tumor growth and mean survival times in both studies were significantly shorter for all non-open field arms (p < 0.05; Log Rank for survival; 2-way ANOVA for tumor growth). Conclusions: Two novel mini-GRIDs were characterized and tested in combination with anti-PD-1 therapy. In this study, neither single dose nor fractionated GRID therapy with anti-PD-1 improved tumor growth delay or survival. Similarly, hemi-irradiation resulted in worse tumor control compared to conventional open field radiotherapy.

## 1. Introduction

Immune checkpoint blockade has revolutionized treatment of multiple cancers through stimulating an anti-tumor immune response largely mediated by cytotoxic T cells (1). Monoclonal antibodies targeting the programmed cell death protein 1 (PD-1)/ programmed cell death ligand 1 (PD-L1) immune checkpoint are now commonly used in cancer therapy to stimulate T cell activity (2). In combination with radiation therapy (RT), anti-PD-1/anti-PD-L1 therapy has shown significant benefit for patients with non-small-cell lung cancer, sarcoma, melanoma, and many other solid tumors (3-9). However, RT dose, fractionation and anti-PD-1 therapy timing are all important variables that may impact therapeutic efficacy, and the optimized configuration is not well known (10). Spatially fractionated radiotherapy may provide a complementary approach to enhance the immunostimulatory effects of RT by preserving a component of infiltrative anti-tumor immune cells within the tumor. Clinically, several case reports have shown promising results. In a patient with metastatic non-small-cell lung cancer, the combination of anti-PD-1 therapy and high-dose LATTICE (a three-dimensional GRID technique) radiotherapy resulted in complete response of a large metastasis measuring 63.2 cc. Notably, other lesions treated palliatively with conventional RT with or without chemotherapy or immunotherapy did not achieve complete response in this patient (11). Similar case reports, typically in palliative settings, have demonstrated complete and partial responses to tumors treated with concurrent or sequential spatially fractionated radiotherapy with immunotherapy (11-16). Despite these promising results, the clinical approach to spatially fractionated radiotherapy has been highly heterogeneous, and mechanisms behind the successes remain largely obscure. There is some consensus however that the immune system plays a key role, and that high dose regions (>10 Gy) are likely necessary for inducing the immunomodulatory effects of spatially fractionated radiotherapy (17, 18).

This work explores the efficacy of pencil mini-beam GRID radiotherapy, a type of spatially fractionated radiotherapy, in combination with anti-PD-1 checkpoint blockade. The GRID treatment provides a heterogeneous dose to tumor using a series of small (<1mm) pencil beamlets in a 2D grid pattern, building on prior work from our team (19). The GRID dose distribution has potential to enhance tumor control by promoting anti-cancer immune function: delivering high radiation doses to antigen-releasing tumor cells in the “peaks” (e.g., 10-20 Gy/fraction) while sparing a component of tumor infiltrating immune cells in the “valleys” (<1-2 Gy/fraction) (20, 21). In combination with immune checkpoint blockade, an amplified immunogenic response is anticipated, although any dependence on GRID hole size/pattern, radiation dose fractionation, and peak/valley doses is not well understood. Anti-tumor immune responses are challenging to predict, with tumors demonstrating heterogeneous baseline immune microenvironment and responses to RT (2). Notably, RT has been shown to both induce and impede anti-tumor immune responses (22). Many immune cells are highly radiosensitive, and cytotoxic T lymphocytes can be damaged by as little as 1-2 Gy (23). Furthermore, RT can induce an immunosuppressive environment by sparing the more radioresistant, immunosuppressive regulatory T cells (Tregs) (24). However, high-dose irradiation has also been shown to “prime” the immune system for anti-tumor effects by increasing antigen presentation and stimulating activation and infiltration of cytotoxic T lymphocytes (25).

In addition to single fraction 15 Gy *in vivo* pencil mini-GRID treatment, this work also explores single fraction hemi-tumor irradiation and whole tumor irradiation with concurrent anti-PD-1 therapy. The efficacy of two GRIDs were investigated, one with 1 mm width and center-to-center spacing, and the other with 254 µm width and 508 µm center-to-center spacing. The smaller GRID was hypothesized to yield the best tumor control due to the combination of better tumor coverage as well as better lymphocyte sparing throughout the tumor than the hemi-field or open field irradiation. Finally, the effect of fractionation was investigated with follow up studies of three fractions of 15 Gy.

## 2. Methods and Materials

### 2.1 GRID design and characterization

In-house mini-GRIDs were designed and constructed from 3mm thick sheets of lead with precision milled drill holes. Holes were equally spaced in a hexagonal formation, as described in prior work (19). All GRID therapy experiments were performed on the XStrahl Small Animal Radiation Research Platform (SARRP) using a 225 kV x-ray beam at 13 mA (26-30). The SARRP contains a rotating gantry, portal fluoroscopy, and a motorized variable collimator. The SARP was fully commissioned for pre-clinical work (see (31) for details) during which the open field output was verified as 3.66Gy/min at 2cm depth in water, 33cm SSD, with a HVL of 0.66 mm Cu. Output measurement was made following TG61 protocol with a farmer chamber and independent verification with EBT3 film. GRID treatment of a mouse leg proceeded with the GRID placed on a support table above the tumor, normal to the incident beam (details in section 2.3). Dosimetric characterization (peak output, peak-to-valley ratio, etc.) was performed with EBT3 film on both GRIDs: a 1 mm beamlet width with 1 mm center-to-center spacing (the 1 mm GRID) and a 254 µm beamlet width with 508 µm center-to-center spacing (the 254 µm GRID). For each, the output factor was determined from the following expression:

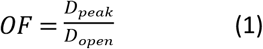

where *OF* is the output factor, *D*_*peak*_ is the average dose delivered in the peak regions, and *D*_*open*_ is the average dose delivered to the open field with no GRID blocking the beam. Doses were determined from EBT3 film following the methods described in (31) utilizing median filter techniques to preserve the peaks and valley doses. EBT3 calibration curves were determined from film exposed to know doses from open field exposures on the SARP from 0-6Gy. All films were scanned 24h post irradiation on an Epsom 11000XL flat-bed scanner. Treatment times were calculated by dividing the time to deliver the prescribed surface dose for the open reference field by the output factor of each GRID.

The peak-to-valley dose ratios (*PVDR*), were determined from the following expression:

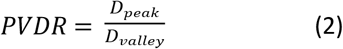

where *D*_*peak*_ is the average maximum dose delivered in the peaks, and *D*_*valley*_ is the dose delivered to the valleys in between beamlets (all doses determined from EBT3 film according to methods outlined in (19). A design goal for the grid was to achieve *D*_*valley*_ <1.5 Gy, a level reported to spare functionality of ∼50% of T cells (32).

### 2.2 Cell Culture

Lewis Lung Carcinoma cells (LLC-1, purchased from Duke University’s Cell Culture Facility) were cultured in Dulbecco’s Modified Eagle’s Medium (Gibco) with 10% v/v fetal bovine serum (ThermoFisher) and 1% v/v penicillin-streptomycin (Gibco). Cells were grown in a humidified environment at 37°C and 5% CO_2_. Prior to transplanting into mice, cells were tested for murine pathogens.

### 2.3 Animal Experiments

All animal studies were performed in accordance with Duke University’s Institutional Animal Care and Use Committee and adhere to the NIH Guide for the Care and Use of Laboratory Animals. C57Bl/6J mice were purchased from Jackson Labs at 6 weeks of age. Tumors were induced in all mice by subcutaneous injection of 1.5 × 10^6^ cells in 100 µL of 50% v/v Matrigel (Corning) and PBS into the flank of each mouse. Mice were monitored thrice weekly, recording weights and bidirectional tumor measurements with calipers. Tumor volume was calculated according to V=(π/6).a.b^2^, where *a* is the length of the longest dimension of the tumor, and *b* is the dimension directly perpendicular to *a*. When all tumors reached >50 mm^3^, mice were randomized into treatment groups. We mitigated any potential bias from initial tumor size by stratifying by tumor size when randomizing mice into groups. Among the trial groups, the mean tumor size at treatment initiation was 290 mm^3^ (SD 17 mm^3^). Care was taken to minimize suffering by daily monitoring and maintenance care, and strict following of humane endpoints for euthanasia including: tumors >15 mm in any dimension or >2000 mm3, weight loss exceeding 15% of free-feeding body weight relative to age-matched control, self mutilation, tumor ulceration, or poor health condition (infection, delayed wound healing, behavioral change, poor posture or ambulating difficulty, lack of grooming, reduced activity level, painful facial expression, signs of moderate to severe pain or distress), or end of study (remaining mice after all mice in GRID treatment groups were euthanized based on reaching tumor size or ulceration endpoint). Isoflurane was administered for anesthesia during radiation therapy, and there was no use of analgesics. Euthanasia was performed using standard CO2 inhalation followed by decapitation.

Two experiments were completed (see Figure 1): the first with the 1 mm GRID and a single fraction of RT, while the second with both the 1 mm and 254 µm GRID and with fractionated RT. In all cases the prescribed dose was delivered to the surface of the tumor, from a single anterior-posterior field. In the first study, mice were randomized into three groups (n=5/group): 1) 15 Gy x 1 fraction, open field; 2) 15 Gy (peak dose) x 1 fraction, 1 mm GRID; and 3) 24 Gy (peak dose) x 1 fraction, 1mm GRID. The higher prescription dose of 24 Gy for the third arm was selected because this is the maximum dose that could be delivered to the peaks while keeping valley dose <1.5 Gy. For the second study, mice were randomized into four groups (n=6-7/group): 1) 15 Gy x 3 fractions, open field (n=6); 2) 15 Gy x 3 fractions, hemi-tumor irradiation (half tumor coverage due to half beam block, n=6); 3) 15 Gy (peak dose) x 3 fractions, 1 mm GRID (n=6); and 4) 15 Gy (peak dose) x 3 fractions, 254 µm GRID. The goal of the fractionation scheme (every 3 days) was to provide a brief window for lymphocyte recovery and action. Daily fractionation is a more common clinical schema; however, there has been some suggestion that fewer fractions of higher dose provide the least decrease in lymphocyte numbers (21). The collimator was set to a 20 × 20 mm field size to ensure that the GRID fully encompassed all tumors with sufficient margin for setup error. In addition, the location of the tumors (subcutaneous flank tumors) was such that peripheral parts of the field laterally were ‘flashing’ outside the animal, limiting the normal tissue volume in the GRID beam.

**Fig 1.**
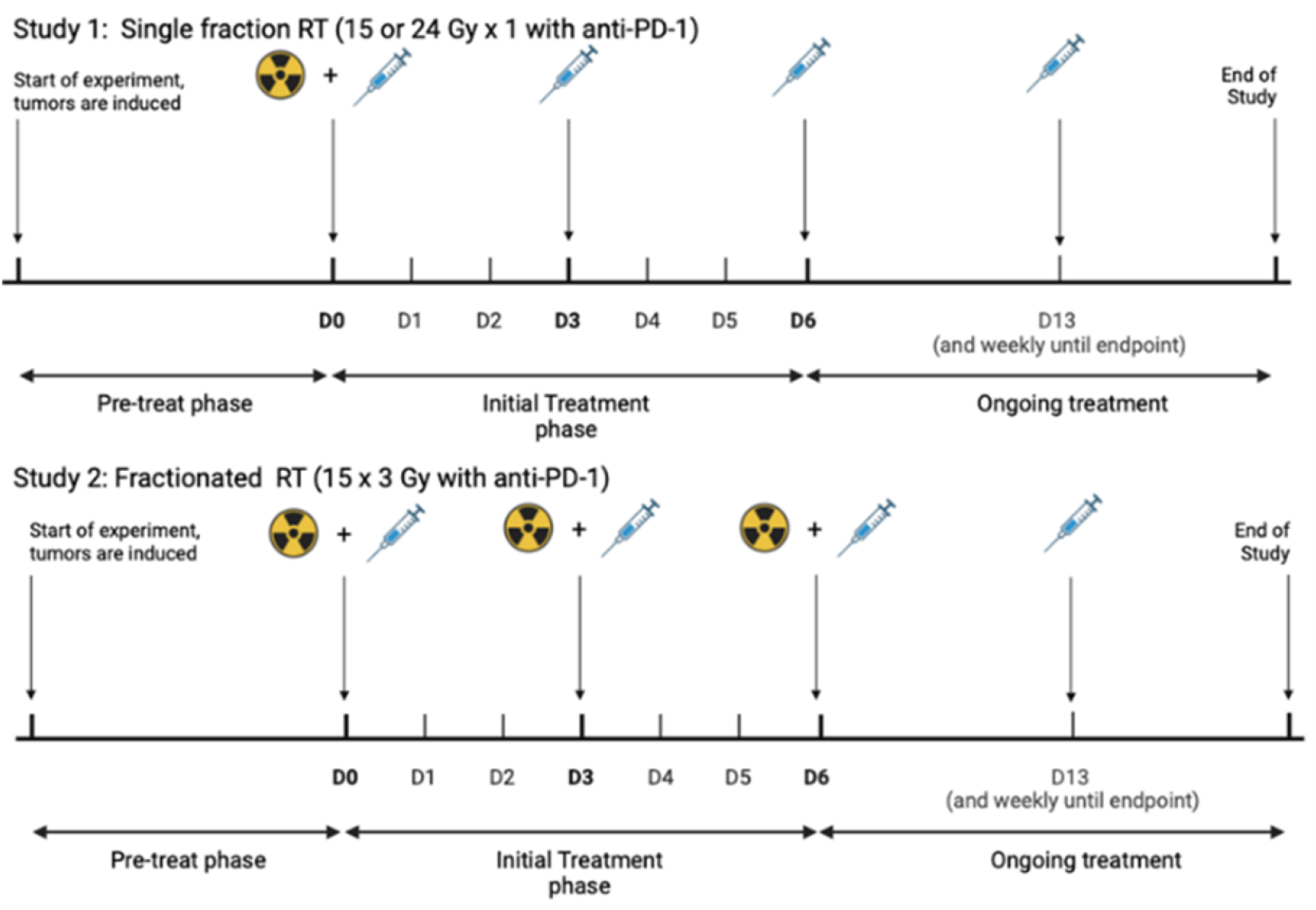
Radiation and anti-PD-1 treatment administration timeline for both single fraction (Study 1) and fractionated (Study 2) radiation experiments. All mice were induced with a LLC-1 flank tumor model. The experimental groups in Study 1 were open field (15 Gy x 1), 1 mm GRID (15 Gy x 1 and 24 Gy x 1 to the GRID peaks). The experimental groups in Study 2 were open field (15 Gy x 3), hemi-tumor irradiation (15 Gy x 3), 1 mm GRID (15 Gy x 3 to peaks) and 254 µm GRID (15 Gy x 3 to peaks).

In addition to RT, all mice received 200 μg of anti-PD-1 antibody (InVivoMAb anti-mouse PD-1, CD279) delivered as an intraperitoneal injection on days 0, 3, and 6 and then weekly thereafter. When both RT and anti-PD-1 coincided on the same day, anti-PD-1 was administered at least 1 hour prior to RT. The anti-PD-1 and RT schedules are shown in Figure 1. Importantly, anti-PD1 alone has been shown in multiple studies to have no significant effect in this model (33-35), therefore our primary control arm was open-field RT + anti-PD1. Both test arms (partial-field + anti-PD1 and GRID field + anti-PD1) would have equal or increased tumor control compared to the primary control if immune activation amplification occured. The treatment configuration and set-up for each mouse is shown in Figure 2. Each GRID was placed on a 3D-printed table <3 cm above the tumor in the SARRP (Figure 2a). Placement of the GRID over the tumor and the treatment field were verified using fluoroscopic imaging for each mouse as shown in Figure 2b.

**Fig 2.**
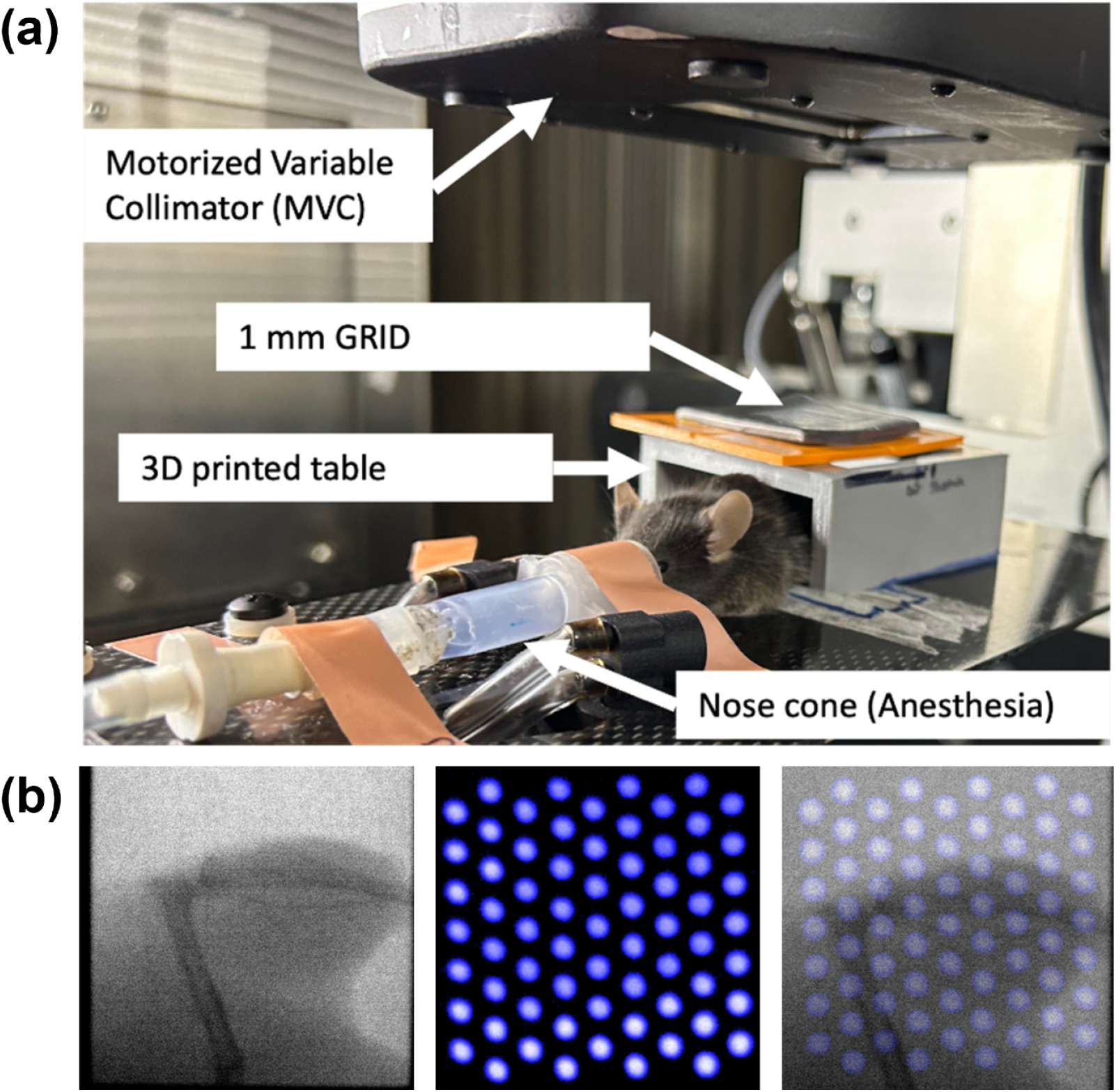
Experimental setup of GRID treatment. (a) In the XStrahl SARRP, the mouse is under isoflurane anesthesia with the tumor-bearing hind limb immobilized directly under the beam. The motorized variable collimator (MVC) set the beam size to be 20 × 20 mm. The 1 mm GRID is placed directly above the tumor on a 3D-printed table. (b) Confirmation of GRID placement was made via portal fluoroscopy. The flank with the tumor is shown on the leftmost panel, followed by the GRID pattern without an underlying tumor. The overlay demonstrates the grid irradiation pattern on the tumor.

### 2.4 Statistical analysis

For tumor growth delay analysis, a 2-way ANOVA with Tukey’s post-hoc test was used to compare experimental groups. Survival was estimated by Kaplan-Meier curves, which were compared with a log-rank test. For all tests, a p-value of <0.05 was considered statistically significant. All tests were performed in GraphPad Prism (v. 10.1.2).

## 3. Results

### 3.1 Characteristics of in-house mini-GRIDs

Figure 3 shows film scans of dose incident on the tumor surface of both the 1mm and 254 µm GRIDs and associated derived parameters. Figure 3a and 3b show EBT3 film scans used to calculate peak and valley doses. Both scans show the pencil beam distribution of each GRID with little qualitative evidence of beam penumbra. Figure 3c summarizes the output factors, peak doses, valley doses and PVDR for both GRIDs, confirming the valley dose remained below the <1.5 Gy design goal to spare lymphocytes.

**Figure 3:**
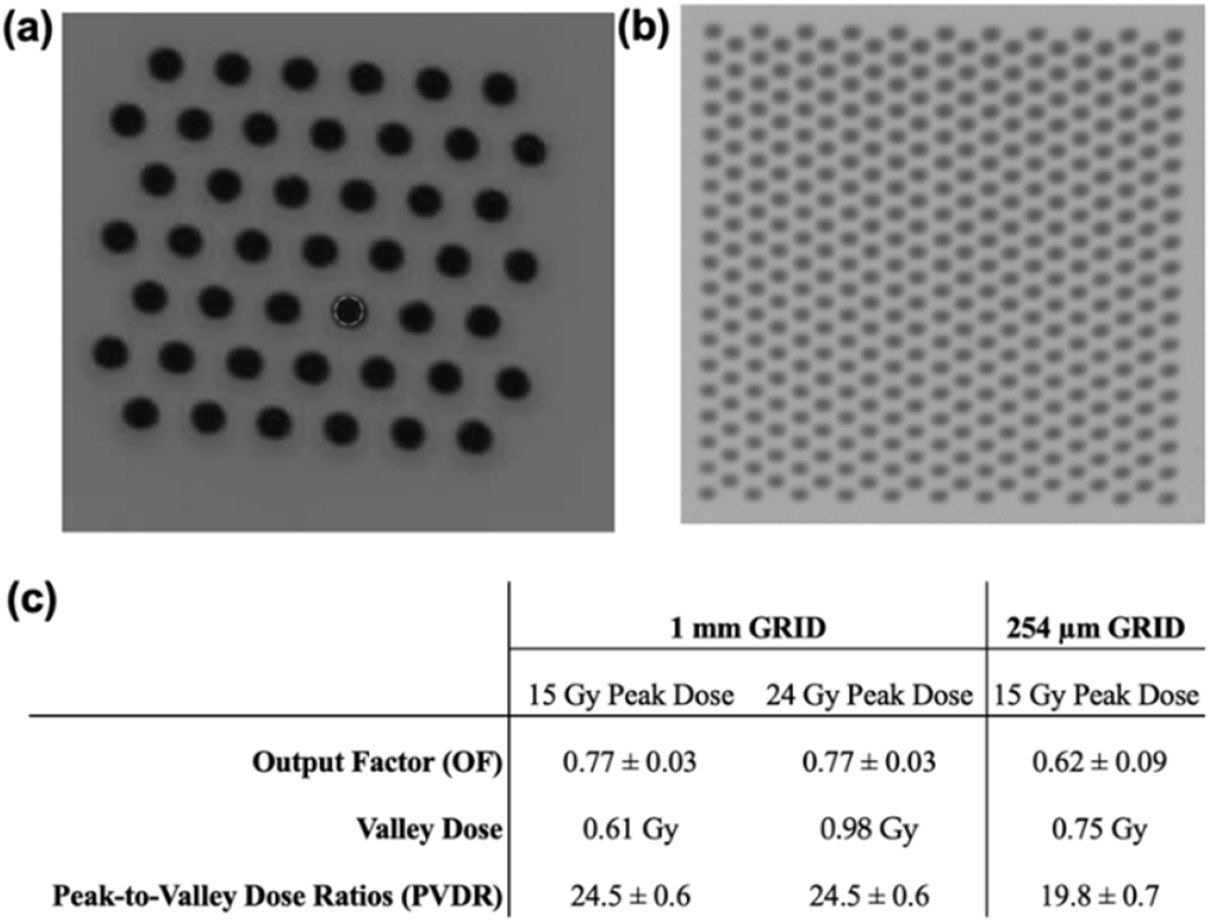
Dosimetric characteristics of GRIDS. (a) The 1 mm GRID and (b) the 254 µm GRID show high resolution GRID patterns in the EBT3 film. (c) The OF and PVDR are reported for each GRID: 1 mm and 254 µm beam diameters, respectively. The average of each is reported ± standard deviation.

### 3.2 Anti-PD-1 immunotherapy combined with GRID RT

Figure 4 shows the tumor growth curves and the survival curves for both the single fraction and fractionated RT studies. For both studies, the open field demonstrated both the longest tumor growth delay, as well as the longest survival. Figure 4a shows a significant difference in tumor growth between the open field and 1 mm GRID – 15 Gy group (p = 0.02, Tukey’s post hoc test), with the latter doubling in tumor volume nearly twice as fast as the open field (mean 4.4 vs 7.1 days, respectively). There is a strong suggestion of tumor growth delay in the higher dose 24 Gy arm as expected, but even the higher GRID dose shows no benefit over the open field treatment at 15 Gy. Similarly, for the multi-fraction study (Figure 4b), the open field group had significantly slower tumor growth compared to all groups (p<0.0001 for all, Tukey’s post hoc test). The benefit of fractionation is clear, with much lower tumor volumes on all days post-RT than corresponding volumes in the single fraction arms.

**Fig 4.**
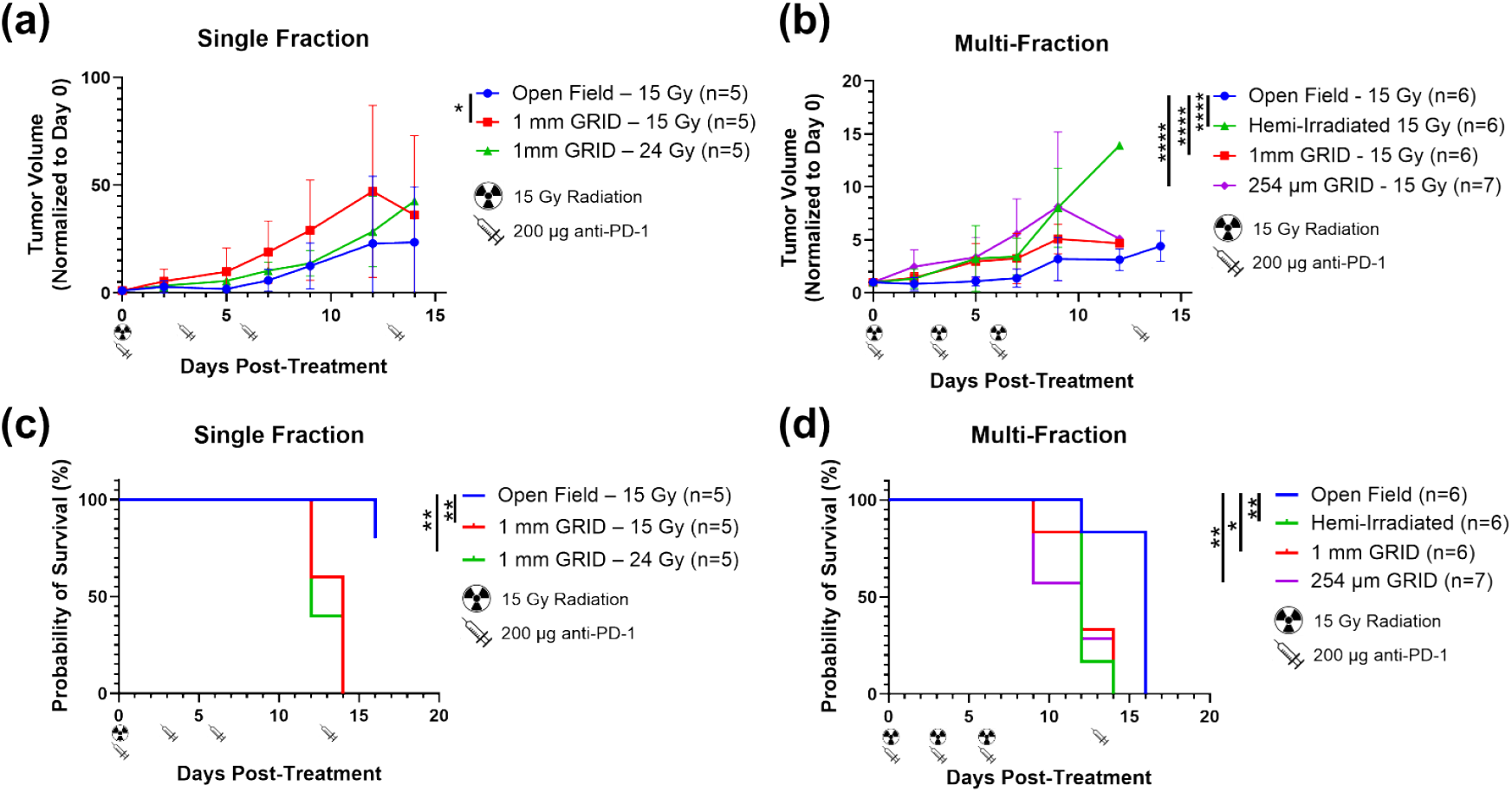
LLC1 flank tumor growth and survival curves for single and multi-fraction GRID treatments. Tumor volumes were normalized to the first day of treatment (day 0) (see schema in Fig 1). Anti-PD1 alone has been shown in multiple studies to have no significant effect in this model (33-35), therefore our primary control arm was open-field RT + anti-PD1. (a) Mice were treated with 15 Gy RT open field, 15 Gy with a 1 mm center-to-center GRID or 24 Gy with a 1 mm center-to-center GRID. Mice were also treated with anti-PD-1 on days 0, 3, and 6 and then weekly thereafter. The open field decreased the tumor growth rate significantly compared to the 15 Gy with 1 mm GRID (p=0.02, Tukey’s post-hoc test). (b) Tumor growth for fractionated treatment with two GRIDs (1 mm or 254 µm). Mice (n=6-7/group) were treated on days 0, 3, and 6 with both anti-PD-1 and 15 Gy radiation/fraction with weekly anti-PD-1 administration thereafter. Mice treated with open field (15 Gy) had a significantly slower tumor growth rate than hemi-irradiation (irradiating half the tumor with a half beam block), the 1 mm GRID, and the 254 µm GRID (p<0.0001 for all, Tukey’s post hoc test). (c) In the single fraction experiment, the open field group had significantly longer survival than the other groups (p = 0.0061, 1 mm GRID – 15 Gy; p = 0.0039, 1 mm GRID – 24 Gy; log-rank). (d) In the fractionated RT experiment, the open field group had significantly longer survival compared to the other groups (p = 0.0046, hemi-irradiation; p = 0.0389, 1 mm GRID log; p = 0.0100, 254 µm GRID; log-rank).

Survival times show similar patterns as tumor growth delay. As shown in Figure 4c, mice treated with open field single fraction RT survived significantly longer than mice treated with 1 mm GRID – 15 Gy (p = 0.0061, log-rank) or 1 mm GRID – 24 Gy (p = 0.0039, log-rank). Similarly, mice treated with fractionated open field RT survived the longest (median 16 days) compared to all groups (p = 0.0046, hemi-irradiated; p = 0.0389, 1 mm GRID; p = 0.0100, 254 µm GRID; log-rank, Figure 4d).

## 4. Discussion

The GRID/anti-PD-1 combination treatment in the studies presented here did not illicit any beneficial effect in terms of either tumor growth delay or survival compared to standard open field treatments of the whole tumor. The addition of anti-PD1 therapy did not achieve the desired goal of amplifying an anti-tumor immune response even in the case of dose escalation in the peaks to 24 Gy, over the 15 Gy open field. No benefit was observed in either the single fraction or fractionated cases, over the whole tumor irradiation arm. A significant benefit was observed in both tumor growth delay and survival in the fractionated cases when compared to the single fraction, but this is to be expected due to increased total dose in these arms. Even for fractionated cases, no benefit to either GRID treatment was observed over the open field irradiation. Tumors in the hemi-irradiated fractionated group grew significantly faster than tumors in the open-field group (p<0.0001), even with concurrent anti-PD-1 therapy. The two GRIDs (1 mm and 254 µm hole size) characterized here approach the size of microbeam GRIDs, which commonly harbor a 50 µm slit-size with 200-400 µm center-to-center spacing (29, 36-39). An advantage of our GRIDs is that they can be used in preclinical cabinet irradiators. At orthovoltage exposures, our GRIDs allow for hypofractionated doses of >15 Gy at the peaks while simultaneously sparing radiosensitive lymphocytes at the valleys (<1.5 Gy). This provides a high-resolution irradiated pattern with small, precise peaks due to the sharp penumbra of a 225 kV beam. We estimate that both GRIDs provide ∼50% tumor-volume coverage. Tumor coverage was defined as the overlap of GRID high dose regions with the mouse tumors. Spatially fractionated RT has been demonstrated to increase CD8+ T cell activity and immune-mediated tumor control in LLC-1 models (40, 41).

It is interesting to contrast our observations with those of Markovsky et al (40), who also compared hemi-irradiation and open-field irradiation with a single 15 Gy treatment in the same flank LLC-1 tumors as used here. They reported hemi-irradiated tumors demonstrated similar tumor growth delay to tumors treated with open field irradiation in immunocompetent mice. It is unclear why their observations differed from those presented here. Differences in microbiome could play a role, as mice of the same strain and age may exhibit different microbiomes based on the housing environment at different institutions, which can affect immune-mediated outcomes (42). However, another research group using LATTICE radiation in an LLC-1 model showed no difference in tumor-growth delay between 50% tumor-volume irradiation and no treatment (41). Fifty percent tumor-volume irradiation using LATTICE targeted the tumor center rather than a lateral half, which may contribute to the lack of tumor response. Notably, the microenvironment is highly variable across the tumor. The tumor periphery is a hub of activity between the tumor and infiltrating immune cells; notably, immature myeloid cells in this region contribute to immune evasion by inhibiting differentiation of dendritic cells, and tumor-associated macrophages contribute to invasiveness and migration (41, 43-45). In addition, the tumor periphery tends to demonstrate higher rates of epithelial-mesenchymal transition, which further promotes tumor invasion and provides a hostile environment for cytotoxic immune cells (46). During hemi-irradiation, only 50% of the tumor periphery is treated, potentially creating a region that is primed for anti-tumor immune responses.

Spatially fractionated radiotherapy has achieved some success clinically in case reports and clinical trials (21, 47, 48). Notably, results may depend on the irradiated tumor volume and pattern of irradiation, and systemic anti-tumor immune effects may come at the expense of local tumor control (41, 49). The variety of GRID patterns and techniques in use makes drawing conclusions on how to optimize spatially fractionated therapy difficult. While this work did not find any benefit, GRID treatments may have benefit in other conditions. For example, stereotactic body radiation therapy with LATTICE (a 3D GRID technique) has risen in clinical use, while microbeam GRIDs are being investigated with synchrotron radiation sources (i.e., FLASH).

## 4. Conclusions

The two mini-GRIDs characterized in this study with 225kV X-rays demonstrate a high PVDR with peaks >15 Gy and valleys <1.5 Gy both to treat the tumor and potentially spare some radiosensitive tumor-infiltrating T cells. The studies presented here, combining GRID therapy with anti-PD-1 therapy, did not find any benefit over conventional whole tumor irradiation, exhibiting generally worse local tumor control with single dose or fractionated radiotherapy compared to an open field irradiating the whole tumor.

## Acknowledgements

We would like to thank the Physics Machine Shop at Duke University for manufacturing the GRIDs used in our experiments.

## References

1. Saxena A. Combining radiation therapy with immune checkpoint blockade for the treatment of small cell lung cancer. Semin Cancer Biol. 2023;90:45–56.

2. Zhang Z, Liu X, Chen D, Yu J. Radiotherapy combined with immunotherapy: the dawn of cancer treatment. Signal Transduct Target Ther. 2022;7(1):258.

3. Chen D, Menon H, Verma V, Guo C, Ramapriyan R, Barsoumian H, et al. Response and outcomes after anti-CTLA4 versus anti-PD1 combined with stereotactic body radiation therapy for metastatic non-small cell lung cancer: retrospective analysis of two single-institution prospective trials. J Immunother Cancer. 2020;8(1).

4. Hiniker SM, Reddy SA, Maecker HT, Subrahmanyam PB, Rosenberg-Hasson Y, Swetter SM, et al. A Prospective Clinical Trial Combining Radiation Therapy With Systemic Immunotherapy in Metastatic Melanoma. Int J Radiat Oncol Biol Phys. 2016;96(3):578–88.

5. Luke JJ, Lemons JM, Karrison TG, Pitroda SP, Melotek JM, Zha Y, et al. Safety and Clinical Activity of Pembrolizumab and Multisite Stereotactic Body Radiotherapy in Patients With Advanced Solid Tumors. J Clin Oncol. 2018;36(16):1611–8.

6. Rodriguez-Ruiz ME, Perez-Gracia JL, Rodriguez I, Alfaro C, Onate C, Perez G, et al. Combined immunotherapy encompassing intratumoral poly-ICLC, dendritic-cell vaccination and radiotherapy in advanced cancer patients. Ann Oncol. 2018;29(5):1312–9.

7. Jiang H, Yu K, Cui Y, Ren X, Li M, Yang C, et al. Combination of Immunotherapy and Radiotherapy for Recurrent Malignant Gliomas: Results From a Prospective Study. Front Immunol. 2021;12:632547.

8. Sahebjam S, Forsyth PA, Tran ND, Arrington JA, Macaulay R, Etame AB, et al. Hypofractionated stereotactic re-irradiation with pembrolizumab and bevacizumab in patients with recurrent high-grade gliomas: results from a phase I study. Neuro Oncol. 2021;23(4):677–86.

9. Lin Z, Cai M, Zhang P, Li G, Liu T, Li X, et al. Phase II, single-arm trial of preoperative short-course radiotherapy followed by chemotherapy and camrelizumab in locally advanced rectal cancer. J Immunother Cancer. 2021;9(11).

10. Boustani J, Lecoester B, Baude J, Latour C, Adotevi O, Mirjolet C, et al. Anti-PD-1/Anti-PD-L1 Drugs and Radiation Therapy: Combinations and Optimization Strategies. Cancers (Basel). 2021;13(19).

11. Jiang L, Li X, Zhang J, Li W, Dong F, Chen C, et al. Combined High-Dose LATTICE Radiation Therapy and Immune Checkpoint Blockade for Advanced Bulky Tumors: The Concept and a Case Report. Front Oncol. 2020;10:548132.

12. Blanco Suarez JM, Amendola BE, Perez N, Amendola M, Wu X. The Use of Lattice Radiation Therapy (LRT) in the Treatment of Bulky Tumors: A Case Report of a Large Metastatic Mixed Mullerian Ovarian Tumor. Cureus. 2015;7(11):e389.

13. Amendola BE, Perez NC, Wu X, Blanco Suarez JM, Lu JJ, Amendola M. Improved outcome of treating locally advanced lung cancer with the use of Lattice Radiotherapy (LRT): A case report. Clin Transl Radiat Oncol. 2018;9:68–71.

14. Amendola BE, Perez NC, Mayr NA, Wu X, Amendola M. Spatially Fractionated Radiation Therapy Using Lattice Radiation in Far-advanced Bulky Cervical Cancer: A Clinical and Molecular Imaging and Outcome Study. Radiat Res. 2020;194(6):724–36.

15. Iori F, Botti A, Ciammella P, Cozzi S, Orlandi M, Iori M, et al. How a very large sarcomatoid lung cancer was efficiently managed with lattice radiation therapy: a case report. Ann Palliat Med. 2022;11(11):3555–61.

16. Ferini G, Castorina P, Valenti V, Illari SI, Sachpazidis I, Castorina L, et al. A Novel Radiotherapeutic Approach to Treat Bulky Metastases Even From Cutaneous Squamous Cell Carcinoma: Its Rationale and a Look at the Reliability of the Linear-Quadratic Model to Explain Its Radiobiological Effects. Front Oncol. 2022;12:809279.

17. Lukas L, Zhang H, Cheng K, Epstein A. Immune Priming with Spatially Fractionated Radiation Therapy. Curr Oncol Rep. 2023;25(12):1483–96.

18. Iori F, Cappelli A, D’Angelo E, Cozzi S, Ghersi SF, De Felice F, et al. Lattice Radiation Therapy in clinical practice: A systematic review. Clin Transl Radiat Oncol. 2023;39:100569.

19. Johnson TR, Bassil AM, Williams NT, Brundage S, Kent CL, Palmer G, et al. An investigation of kV mini-GRID spatially fractionated radiation therapy: dosimetry and preclinical trial. Phys Med Biol. 2022;67(4).

20. Sammer M, Teiluf K, Girst S, Greubel C, Reindl J, Ilicic K, et al. Beam size limit for pencil minibeam radiotherapy determined from side effects in an in-vivo mouse ear model. PLoS One. 2019;14(9):e0221454.

21. Yan W, Khan MK, Wu X, Simone CB, 2nd, Fan J, Gressen E, et al. Spatially fractionated radiation therapy: History, present and the future. Clin Transl Radiat Oncol. 2020;20:30–8.

22. Park B, Yee C, Lee KM. The effect of radiation on the immune response to cancers. Int J Mol Sci. 2014;15(1):927–43.

23. Heylmann D, Ponath V, Kindler T, Kaina B. Comparison of DNA repair and radiosensitivity of different blood cell populations. Sci Rep. 2021;11(1):2478.

24. Liu S, Sun X, Luo J, Zhu H, Yang X, Guo Q, et al. Effects of radiation on T regulatory cells in normal states and cancer: mechanisms and clinical implications. Am J Cancer Res. 2015;5(11):3276–85.

25. Gupta A, Probst HC, Vuong V, Landshammer A, Muth S, Yagita H, et al. Radiotherapy promotes tumor-specific effector CD8+ T cells via dendritic cell activation. J Immunol. 2012;189(2):558–66.

26. Bache ST, Juang T, Belley MD, Koontz BF, Adamovics J, Yoshizumi TT, et al. Investigating the accuracy of microstereotactic-body-radiotherapy utilizing anatomically accurate 3D printed rodent-morphic dosimeters. Med Phys. 2015;42(2):846–55.

27. Trappetti V, Potez M, Fernandez-Palomo C, Volarevic V, Shintani N, Pellicioli P, et al. Microbeam Radiation Therapy Controls Local Growth of Radioresistant Melanoma and Treats Out-of-Field Locoregional Metastasis. Int J Radiat Oncol Biol Phys. 2022;114(3):478–93.

28. Cramer CK, Yoon SW, Reinsvold M, Joo KM, Norris H, Hood RC, et al. Treatment Planning and Delivery of Whole Brain Irradiation with Hippocampal Avoidance in Rats. PLoS One. 2015;10(12):e0143208.

29. Yoon SW, Kodra J, Miles DA, Kirsch DG, Oldham M. A method for generating intensity-modulated radiation therapy fields for small animal irradiators utilizing 3D-printed compensator molds. Med Phys. 2020;47(9):4363–71.

30. Rankine LJ, Newton J, Bache ST, Das SK, Adamovics J, Kirsch DG, et al. Investigating end-to-end accuracy of image guided radiation treatment delivery using a micro-irradiator. Phys Med Biol. 2013;58(21):7791–801.

31. Brundage S. Commissioning a State-of-Art Small Animal Irradiator and Novel Mini-GRID Treatment Technique [Master’s Thesis]. https://hdl.handle.net/10161/26886: Duke University; 2022.

32. Nakamura N, Kusunoki Y, Akiyama M. Radiosensitivity of CD4 or CD8 positive human T-lymphocytes by an in vitro colony formation assay. Radiat Res. 1990;123(2):224–7.

33. Xia L, Tian E, Yu M, Liu C, Shen L, Huang Y, et al. RORgammat agonist enhances anti-PD-1 therapy by promoting monocyte-derived dendritic cells through CXCL10 in cancers. J Exp Clin Cancer Res. 2022;41(1):155.

34. Ajona D, Ortiz-Espinosa S, Lozano T, Exposito F, Calvo A, Valencia K, et al. Short-term starvation reduces IGF-1 levels to sensitize lung tumors to PD-1 immune checkpoint blockade. Nat Cancer. 2020;1(1):75–85.

35. Zilionyte K, Bagdzeviciute U, Mlynska A, Urbstaite E, Paberale E, Dobrovolskiene N, et al. Correction to: Functional antigen processing and presentation mechanism as a prerequisite factor of response to treatment with dendritic cell vaccines and anti-PD-1 in preclinical murine LLC1 and GL261 tumor models. Cancer Immunol Immunother. 2022;71(11):2701.

36. Potez M, Bouchet A, Flaender M, Rome C, Collomb N, Grotzer M, et al. Synchrotron X-Ray Boost Delivered by Microbeam Radiation Therapy After Conventional X-Ray Therapy Fractionated in Time Improves F98 Glioma Control. Int J Radiat Oncol Biol Phys. 2020;107(2):360–9.

37. Adam JF, Balosso J, Bayat S, Berkvens P, Berruyer G, Brauer-Krisch E, et al. Toward Neuro-Oncologic Clinical Trials of High-Dose-Rate Synchrotron Microbeam Radiation Therapy: First Treatment of a Spontaneous Canine Brain Tumor. Int J Radiat Oncol Biol Phys. 2022;113(5):967–73.

38. Schultke E, Bayat S, Bartzsch S, Brauer-Krisch E, Djonov V, Fiedler S, et al. A Mouse Model for Microbeam Radiation Therapy of the Lung. Int J Radiat Oncol Biol Phys. 2021;110(2):521–5.

39. Fernandez-Palomo C, Schultke E, Brauer-Krisch E, Laissue JA, Blattmann H, Seymour C, et al. Investigation of Abscopal and Bystander Effects in Immunocompromised Mice After Exposure to Pencilbeam and Microbeam Synchrotron Radiation. Health Phys. 2016;111(2):149–59.

40. Markovsky E, Budhu S, Samstein RM, Li H, Russell J, Zhang Z, et al. An Antitumor Immune Response Is Evoked by Partial-Volume Single-Dose Radiation in 2 Murine Models. Int J Radiat Oncol Biol Phys. 2019;103(3):697–708.

41. Kanagavelu S, Gupta S, Wu X, Philip S, Wattenberg MM, Hodge JW, et al. In vivo effects of lattice radiation therapy on local and distant lung cancer: potential role of immunomodulation. Radiat Res. 2014;182(2):149–62.

42. Forster SC, Clare S, Beresford-Jones BS, Harcourt K, Notley G, Stares MD, et al. Identification of gut microbial species linked with disease variability in a widely used mouse model of colitis. Nat Microbiol. 2022;7(4):590–9.

43. Wyckoff JB, Wang Y, Lin EY, Li JF, Goswami S, Stanley ER, et al. Direct visualization of macrophage-assisted tumor cell intravasation in mammary tumors. Cancer Res. 2007;67(6):2649–56.

44. Gabrilovich DI, Ostrand-Rosenberg S, Bronte V. Coordinated regulation of myeloid cells by tumours. Nat Rev Immunol. 2012;12(4):253–68.

45. Condeelis J, Pollard JW. Macrophages: obligate partners for tumor cell migration, invasion, and metastasis. Cell. 2006;124(2):263–6.

46. Quail DF, Joyce JA. Microenvironmental regulation of tumor progression and metastasis. Nat Med. 2013;19(11):1423–37.

47. Snider JW, Molitoris J, Shyu S, Diwanji T, Rice S, Kowalski E, et al. Spatially Fractionated Radiotherapy (GRID) Prior to Standard Neoadjuvant Conventionally Fractionated Radiotherapy for Bulky, High-Risk Soft Tissue and Osteosarcomas: Feasibility, Safety, and Promising Pathologic Response Rates. Radiat Res. 2020;194(6):707–14.

48. Iturri L, Jouglar E, Gilbert C, Espenon J, Juchaux M, Prezado Y. A first evaluation of the efficacy of minibeam radiation therapy combined with an immune check point inhibitor in a model of glioma-bearing rats. Clin Transl Radiat Oncol. 2025;51:100911.

49. Billena C, Khan AJ. A Current Review of Spatial Fractionation: Back to the Future? Int J Radiat Oncol Biol Phys. 2019;104(1):177–87.

